# Examining the relationship between prenatal depression, amygdala-prefrontal structural connectivity and behaviour in preschool children

**DOI:** 10.1101/692335

**Authors:** Rebecca E. Hay, Jess E. Reynolds, Melody Grohs, Dmitrii Paniukov, Gerald F. Giesbrecht, Nicole Letourneau, Deborah Dewey, Catherine Lebel

## Abstract

Prenatal depression is a common, underrecognized, and undertreated condition with negative consequences on child behaviour and brain development. Neurological dysfunction of the amygdala, cingulate cortex and hippocampus are associated with the development of depression and stress disorders in youth and adults. Although prenatal depression is associated with both child behaviour and neurological dysfunction, the relationship between these variables remains unclear. In this study, fifty-four mothers completed the Edinburgh Depression Scale (EDS) during the second and third trimester of pregnancy and 3 months postpartum. Their children’s behaviour was assessed using the Child Behaviour Checklist (CBCL), and the children had diffusion magnetic resonance imaging (MRI) at age 4.1 +/− 0.8 years. Associations between prenatal depressive symptoms, child behaviour, and child brain structure were investigated. Third trimester EDS scores were associated with altered white matter in the amygdala-frontal tract and the cingulum, controlling for postpartum depression. Externalizing behaviour was sexually differentiated in the amygdala-frontal pathway. Altered structural connectivity between the amygdala and frontal cortex mediated the relationship between third trimester maternal depressive symptoms and child externalizing behaviour in males, but not females. These findings suggest that altered brain structure is a possible mechanism via which prenatal depressive symptoms can impact child behaviour, highlighting the importance of both recognition and intervention in prenatal depression.

## Introduction

Depressive symptoms affect between 13-40% of women in pregnancy [1]. Despite this, prenatal depression is under-recognized and under-treated [2,3], which can lead to poor child outcomes. Independent of postnatal depression and anxiety, prenatal depression is associated with lower child intelligence [4,5], higher infant generalized anxiety and sleep problems [6], increased internalizing and externalizing behaviour in preschool-aged children [7], and increased risk for depression at 18 years [8]. Furthermore, evidence indicates that effective treatment of maternal prenatal depression improves child outcomes, for example, by decreasing internalizing behaviour [9].

Child behavioural deficits associated with prenatal maternal depression might be explained, at least in part, by alterations to the underlying neurological structure and function. The stress response is regulated by the cingulate cortex, hippocampus, and amygdala [10]. Dysfunction of these brain regions is associated with the development of depression and stress disorders in youth and adults [11]. The amygdala is particularly important for emotional processing [12,13], specifically through its connections to the frontal cortex [14]. Altered connectivity between the amygdala and associated cortical areas is associated with vulnerability to depression [15–18], anxiety, and increased stress reactivity [19,20]. Maternal prenatal depression may impact child behaviour by altering this neurological circuit in utero. Indeed, prenatal depression is associated with altered amygdala functional connectivity in 6-month-old infants [21], altered right amygdala microstructure at birth [22], lower frontal and parietal brain activation [7], and significant cortical thinning in the right frontal lobe of children [23,24].

Together, these findings suggest that maternal prenatal depression affects child behaviour by impacting neurological development. However, to our knowledge, no study has investigated whether alterations in brain structure mediate the relationship between prenatal depression and child behaviour, which would provide more concrete evidence of a mechanism. The goal of this study was to determine the relationship between prenatal depression and child brain and behaviour outcomes and investigate whether maternal depression and child behaviour is mediated by alterations to structural connectivity.

## 2. Methods

### 2.1. Participants

Fifty-four mothers and their children (24 female; 4.1 +/− 0.8 years at the MRI scan) were enrolled from an ongoing, prospective study examining maternal nutrition and child outcomes [25]. All participants provided informed consent prior to being included in the study, and this project was approved by the University of Calgary Health Research Ethics Board and the University of Alberta Health Research Ethics Biomedical Panel. Demographics of the women and children are summarized in Table 1. This sample partially overlaps with a sample of children examined in a previous study of brain structure and maternal depression symptoms [24]. Maternal years of postsecondary education (all mothers had completed high school) was used as a proxy for socioeconomic status. Exclusion criteria for children included diagnosed neurological disorders, history of head trauma, genetic disorders impacting cognitive or motor function, and contraindications to MRI.

**Table 1:** Demographic characteristics of mothers and children enrolled in the study. N=54 unless otherwise specified.

| | Range | Mean (± SD) |
| --- | --- | --- |
| **Mothers** |  |  |
| Maternal age at child's birth (years) | 26-38 | 32.3 ± 2.8 |
| Postsecondary education (years) | 0-12 | 5.53 ± 2.8 |
| EDS 1<sup>st</sup> Trimester (n=27) | 0-16 | 4.85 ± 3.7 |
| EDS 2<sup>nd</sup> Trimester (n=47) | 0-16 | 4.47 ± 4.0 |
| EDS 3<sup>rd</sup> Trimester (n=53) | 0-18 | 4.83 ± 3.5 |
| EDS 3 months postpartum | 0-19 | 4.72 ± 4.7 |
| **Children** |  |  |
| Sex | 24 female/30 male |  |
| Gestational age at birth (weeks) | 35-41.86 | 39.3 ± 1.4 |
| Birth weight (g) | 2230-4150 | 3344.9 ± 451.8 |
| Age at scan (years) | 2.85-6.00 | 4.12 ± 0.8 |
| CBCL Externalizing Behaviour T-Scores | 28-68 | 44.81 ± 9.4 |
| CBCL Internalizing Behaviour T-Scores | 20-69 | 44.80 ± 9.2 |

### 2.2. Depressive Symptoms

Maternal depression symptoms were assessed using the Edinburgh Depression Scale (EDS), a 10-item self-report validated scale for measuring depressive symptoms prenatally and postnatally [26,27]. Higher scores indicate worse depressive symptoms; scores ≥12 are usually consistent with a physician diagnosis of major depressive disorder (MDD) [26]. Women completed the EDS once during each trimester of pregnancy (first trimester: 11.0 ± 2.8 weeks *n* = 27; second trimester: 16.8 ± 2.2 weeks, *n* = 47; third trimester 31.5 ± 1.1 weeks, *n* = 53) and at 3 months postpartum (11.5 ± 2.8 weeks, *n* = 54); see Table 1. Due to late enrollment, 27 women did not complete EDS in the first trimester; these scores were not used for analysis. 7 women were missing second trimester scores, and 1 woman was missing a third trimester score. Two women (7.4%) scored ≥12 on the EDS in the first trimester, 4 women (8.5%) in the second trimester, 2 (3.8%) in the third trimester, and 5 (9.3%) postpartum. One participant was taking an antidepressant prenatally (selective serotonin reuptake inhibitor class, moderate dose).

### 2.3. Behavioural Measures

Within 6 months of their MRI scan, each child’s parent completed the Child Behaviour Checklist (CBCL) [28]. The CBCL is a 118-item measure of child behaviour in broad categories including internalizing and externalizing behaviour. Internalizing behaviour incorporates the Anxious/Depressed, Withdrawn, Emotionally-Reactive, and Somatic Complaints domains into a final T-score. Externalizing behaviour incorporates the domains of Attention Problems and Aggressive Behaviour into a final T-score (see Table 1). The CBCL is a useful screening tool for the likelihood of developing a psychiatric condition later in life [29]. T-scores ≥70 are considered in the clinically significant range, and children with T-scores ≥60 considered to be “at-risk”.

### 2.4. Imaging

MRI data was collected at the Alberta Children Hospital on a GE 3T MR750w (General Electric, Waukesha, Wisconsin) using a 32-channel head coil. During the scan, children were either sleeping naturally or watching a movie. Whole brain diffusion tension imaging (DTI) was collected using single shot spin echo echo-planar imaging with 30 diffusion encoding gradient directions at b=750 s/mm^2^ and 5 images at b=0 s/mm^2^, TR=6750 ms, TE=79 ms, and spatial resolution of 1.6×1.6×2.2 mm^3^ (resampled on scanner to 0.78×0.78×2.2 mm).

### 2.5. Image Processing and Analysis

Images were manually checked and volumes with artifacts or motion corruption were removed. All datasets included had ≥18 high quality diffusion weighted and ≥2 b0 volumes. Standard pre-processing of DTI images was performed using ExploreDTI [30], and included correction for signal drift, Gibbs ringing (non-DWIs), eddy currents and motion [31]. Semi-automated deterministic streamline tractography [32] was used to delineate the fornix, cingulum, and white matter connectivity from the amygdala to the prefrontal cortex [33] (Figure 1). The angle threshold was set to 30°, and the minimum fractional anisotropy (FA) threshold was set to 0.20 to initiate and continue tracking. A representative scan (a 3.68 year old female) was identified using FSL [34] for use as a target scan in semi-automated tractography. All other subjects’ FA maps were registered to this template. Regions of interest (ROIs) were drawn in the left and right hemispheres on the template and warped to fit each subject using the inverse registration parameters calculated from the previous step [35]. ROIs for the fornix and cingulum were drawn using a priori information on tract location [32,35,36].

**Figure 1.**
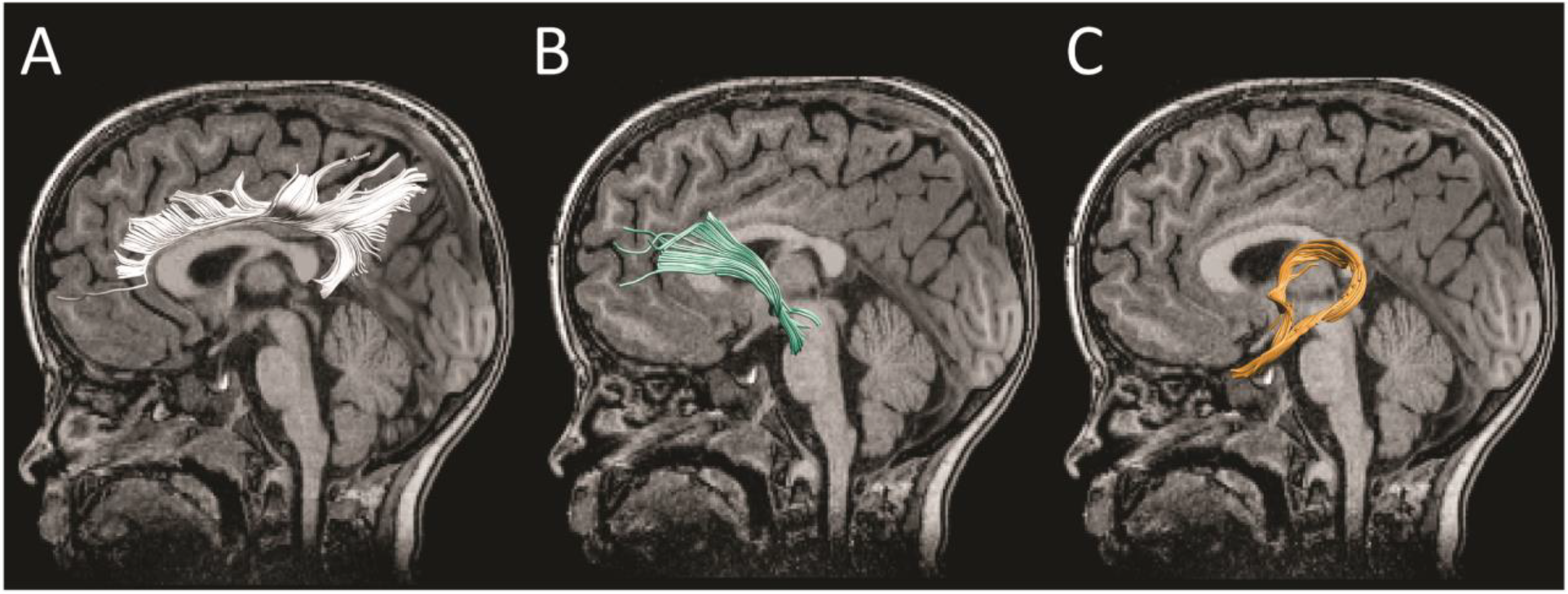
White matter tracts evaluated in this study were the cingulum (A), amygdala pathway (B), and fornix (C)

The amygdala is functionally connected with the anterior cingulate, orbitofrontal, and prefrontal cortex (PFC) [37], subcortical regions including the hippocampus, and projects posteriorly to the occipital cortex, basal ganglia, midbrain, and cerebellum [21]. To delineate the amygdala tract, ROIs were drawn in the axial plane on the amygdala and the coronal plane in the frontal lobe to capture anterior projections [33]. An exclusion region was drawn proximal to the inclusion ROIs through the corona radiata and central sulcus on the same axial slice to isolate the tract of interest. After semi-automated tractography, minor manual edits were done to remove spurious fibers in approximately 15% of the participants. The resulting tract contained fibers emanating from the amygdala with anterior projections to the PFC and posterior projections to the brainstem (Figure 1b). Anterior fibers bundled in the thalamus and projected to the frontal cortex, including the ventral medial prefrontal cortex (vmPFC) and anterior cingulate cortex. The operator was blind to maternal depressive symptoms and child sex and behaviour. Mean FA and mean diffusivity (MD) were extracted for each tract. Left and right tracts were examined separately for the amygdala tract and the cingulum; the fornix was analyzed as a whole.

### 2.6. Regression Analysis

Statistical analysis was performed in IBM SPSS 24. Brain measures (FA and MD for 3 white matter tracts in each hemisphere) were tested separately as dependent variables using linear regression, with age, sex, prenatal trimester EDS, EDS-sex interaction, maternal post-secondary education, gestational age at birth, birthweight, and postnatal EDS as predictors. Second and third trimester EDS symptoms were tested separately. Similarly, brain measures were tested for relationships with behaviour using linear regression with age, sex, CBCL scores (externalizing or internalizing), CBCL-sex interaction, maternal post-secondary education, gestational age at birth, and birthweight as predictors. If the sex interaction terms were not significant, they were removed from the model and the regression was re-run. First trimester EDS symptoms were not tested due to the low number of participants.

Results are reported both uncorrected and corrected for 5 multiple comparisons using false discovery rate (FDR).

### 2.7. Mediation Analysis

White mater measures that were related to both CBCL and EDS scores were selected for a mediation analysis. For white matter measures with significant sex interactions, males and females were analyzed separately. Maternal depressive symptoms (EDS) were entered as the independent variable (X), brain measures as the mediator (M), and CBCL scores as the outcome variable (Y). Non-parametric mediation tests were run using percentile bootstrapping, given the validation of this model on the non-normal distribution of mediating variables [38]. These tests were performed using in-house python scripts.

## 3. Results

### 3.1. Maternal depressive symptoms and white matter measures

Second trimester maternal EDS scores were negatively associated with diffusion parameters in the cingulum. EDS had a main effect on FA values in the left cingulum (F = 4.805, p = 0.035), such that children of women with higher depressive symptoms tended to have lower FA. The EDS-sex interaction was significant for left cingulum MD (F = 4.091, p = 0.050; figure 2 A & B), such that higher maternal depressive symptoms were associated with higher MD in boys and lower MD in girls. These results did not survive FDR multiple comparison correction.

**Figure 2.**
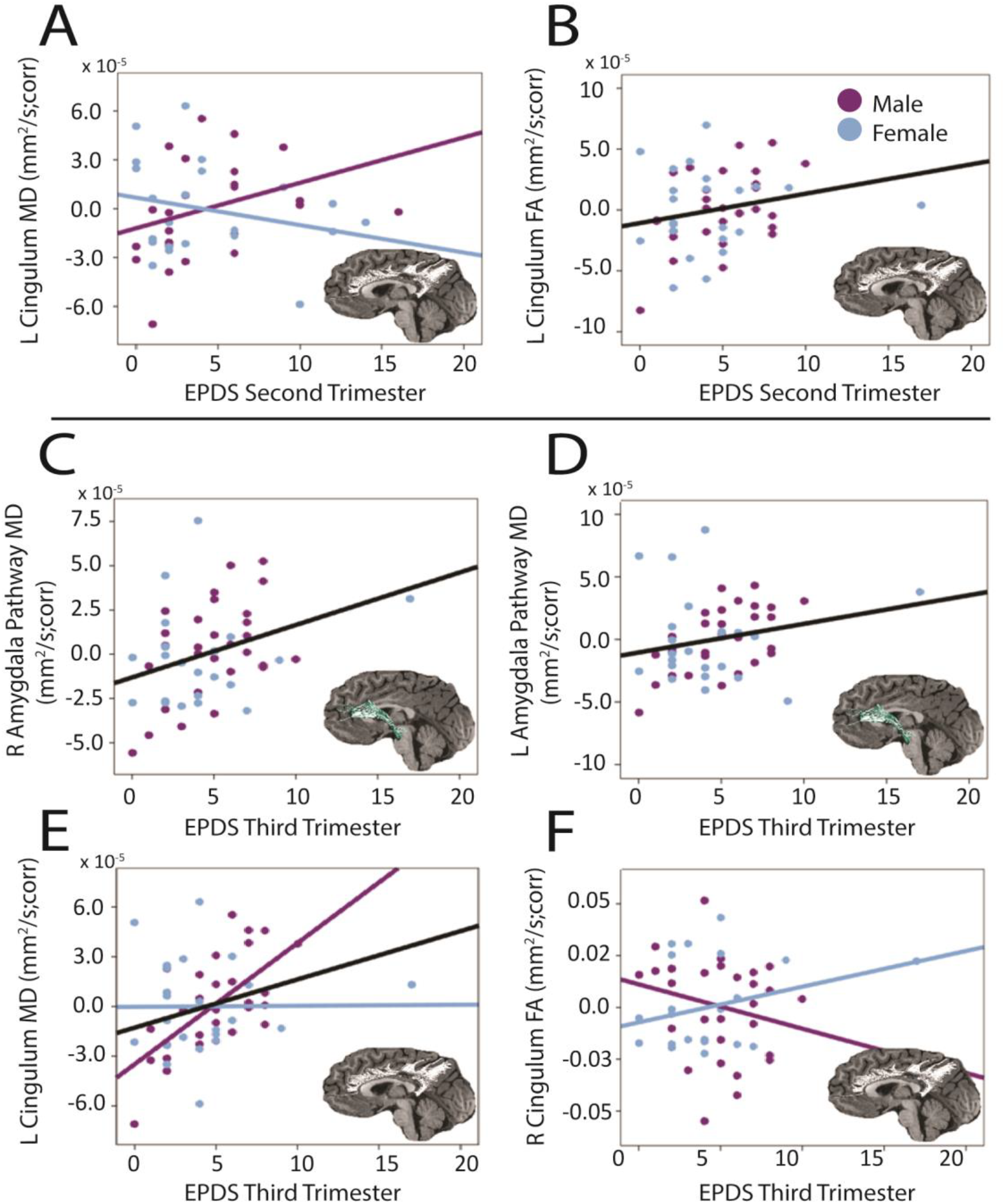
Relationships between white matter measures and maternal depressive symptoms. Black line indicates a main effect of EDS; lines color coded by sex indicate an interaction between EDS*sex. Maternal depressive symptoms in the second trimester were significantly associated with bilateral cingulum MD (A, B). Maternal depressive symptoms in the third trimester were positively associated with the right and left amygdala pathway MD (C,D), the left cingulum MD (E), and were sexually differentiated in the right cingulum FA (F).

Third trimester maternal depressive symptoms were significantly associated with diffusion parameters in the amygdala pathway. EDS had a positive main effect on both right amygdala pathway MD (F = 9.900, p = 0.002), and left amygdala pathway MD (F = 4.520, p = 0.032), as in children of women with higher 3^rd^ trimester depressive symptoms tended to have higher MD. (Figure 2 C & D)

Third trimester EDS had a main effect on MD values in both the left (F = 8.815, p = 0.005) and right cingulum (F = 4.377, p = 0.035), such that mothers with higher third trimester depressive symptoms had children with higher MD in the cingulum bilaterally. (Figure 2 E & F)

Effects of maternal depressive symptoms in the third trimester were sexually differentiated for the cingulum, with a significant sex-EDS interaction in the right cingulum FA (F= 4.403, p = 0.042) and left cingulum MD (F = 5.757 p = 0.021) (Figure 2 E & F) such that higher maternal depressive symptoms were associated with lower FA or higher MD in males. All results for EDS-MD relationships in the amygdala and cingulum survived FDR correction for multiple comparisons. The right cingulum FA interaction did not survive FDR correction.

### 3.2. White matter and child behaviour

The sex-externalizing behaviour interaction was significant for the right amygdala pathway for both FA (F = 7.303, p = 0.010) and MD (F = 6.851 p = 0.012) (Figure 3) and the left amygdala pathway for FA (F = 6.340, p = 0.016) and MD (F = 9.177, p = 0.004) (Figure 3). In all cases, higher externalizing behavioural symptoms were associated with higher FA or lower MD in females, and lower FA or higher MD in males. All results survived FDR correction for multiple comparisons.

**Figure 3.**
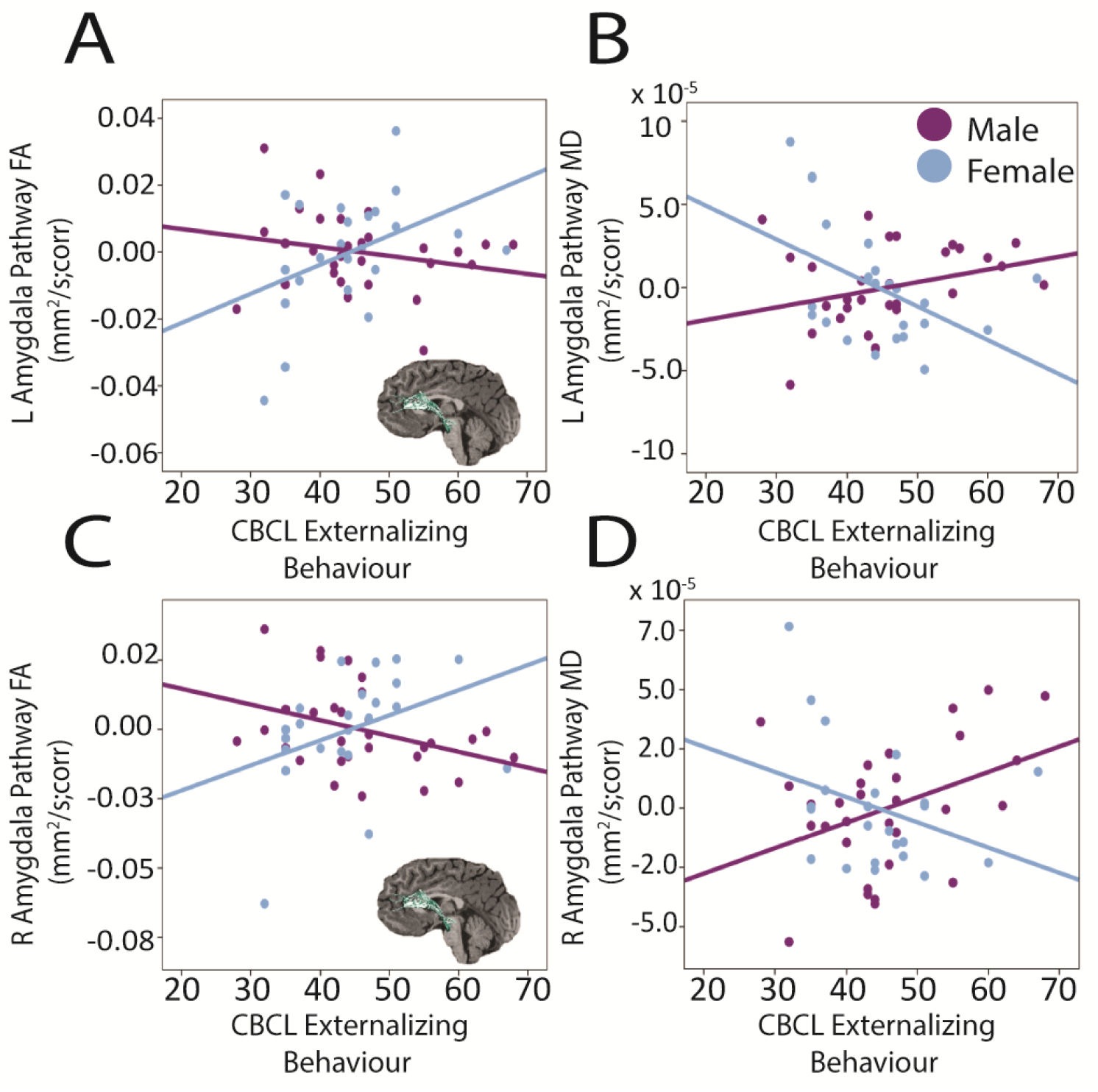
Behavioural measures and child white matter microstructure. Externalizing behaviour was sexually differentiated in the left amygdala pathway (A, B) and right amygdala pathway (C,D).

Internalizing behaviour in children was positively associated with left cingulum FA (F = 5.249, p = 0.036), but did not survive FDR correction.

### 3.3. Mediation analysis

MD of the right amygdala significantly mediated the relationship between third trimester depressive symptoms and externalizing behaviour in males (95% confidence interval: [0.013, 2.05] Figure 4).

**Figure 4.**
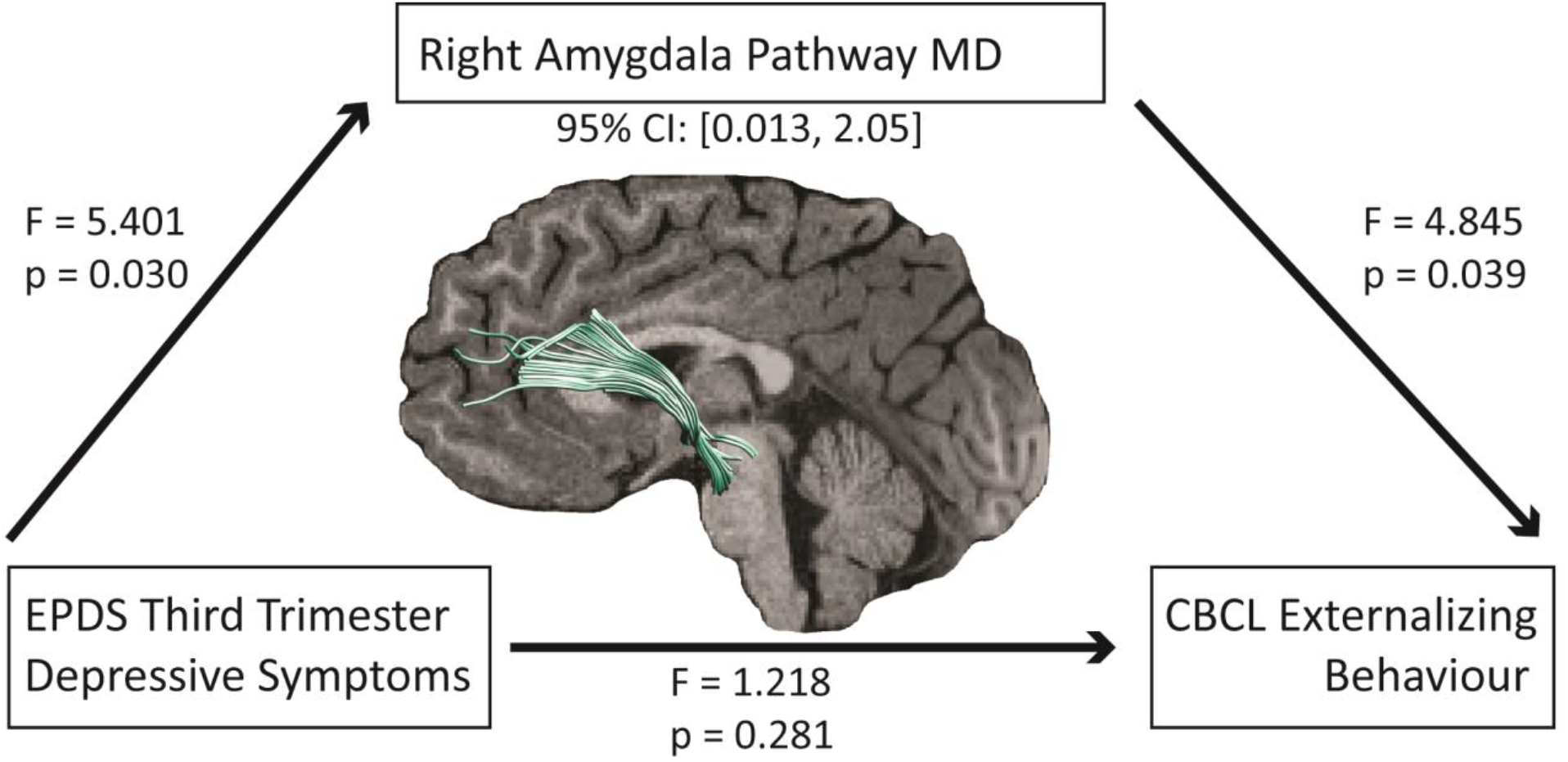
Mean diffusivity of the white matter projections from the right amygdala significantly mediated the relationship between third trimester depressive symptoms and externalizing behaviour in males. The relationship between the right amygdala pathway, third trimester depressive symptoms, and externalizing behaviour were significant. The direct relationship between externalizing behaviour and third trimester depressive symptoms was not significant, but the indirect (mediation) relationship was (95% confidence interval: [0.013, 2.05]). All imaging parameters are corrected for maternal post-secondary education, child age at scan, gestational age, birth weight, and postpartum depressive symptoms.

## 4. Discussion

Here, we show that prenatal maternal depressive symptoms are associated with altered limbic-prefrontal connectivity in young children. Furthermore, reduced structural connectivity between the amygdala and anterior cingulate mediated the relationship between maternal depressive symptoms and externalizing behaviour in boys. These results suggest that reduced brain connectivity within the stress network is a possible mechanism via which maternal depression symptoms impacts children’s behaviour.

We found weaker structural connectivity (lower FA and higher MD) in the amygdala pathway and cingulum associated with higher prenatal maternal depressive symptoms. Because FA increases and MD decreases with age [39], lower FA and/or higher MD values may indicate underdeveloped white matter structures in young children prenatally exposed to higher depression symptoms, which may in turn predispose these children to dysregulated emotional states. Previous functional imaging studies have reported weaker amygdala-PFC functional connectivity in children born to more depressed mothers [22]. Similarly, reduced functional connectivity from the amygdala to the PFC was observed in 4-7 year old children exposed to early life stress such (parental divorce or familial death), and this weaker connectivity was also associated with more difficulties in aggression and attention [40]. On the other hand, Qiu et al (2015) found greater functional connectivity from the left amygdala to frontal cortices in neonates exposed to maternal prenatal depression [21], indicating a potentially complex picture that depends on factors such as a child’s age and sex. Structurally, previous research has found decreased FA in the right amygdala of infants born to mothers with higher third trimester depressive symptoms [41].

The cingulum was also associated with prenatal depressive symptoms, and is thought to be a stress susceptible structure [42]. Lower FA has been associated with dissociation and depression in young adults exposed to early life and prenatal stress [43,44]. The hippocampus receives serotonergic transmission from the midbrain raphe via the fornix and the cingulum [45], providing a potential means by which alterations in the cingulum can disrupt mood.

Previous results from our lab, in an overlapping sample, suggested an accelerated pattern of development (lower diffusivity) in right frontal-temporal white matter in children exposed to greater second trimester depressive symptoms [24]. Indeed, other data has found that potentially accelerated development may predispose children to psychiatric conditions later in life [46–48]. This may be a product of timing and brain areas. In fact, in this study we found a non-significant but similar pattern of accelerated development, where second trimester maternal depressive symptoms positively associated with left cingulum FA in females. It is likely that timing of prenatal maternal depressive symptoms is a key factor in how brain development is affected. Furthermore, it is likely that prenatal depressive symptoms impact areas of the brain differently, as our previous findings were in a large area of right frontal-temporal white matter, while the current results are limited to the cingulum and amygdala tract. Nonetheless, these results paint a complex picture of prenatal environment and the child brain, where outcomes vary by timing of exposure, child age, and sex.

Altered structural connectivity similar to that observed here is associated with depression. Decreased connectivity between the prefrontal cortex and the amygdala is seen in adolescents with MDD [49] and individuals with a familial history of depression [50]. Decreased functional and structural connectivity between the amygdala and frontal cortex may indicate a loss of inhibitory control and subsequent increased amygdala reactivity. This idea is supported by data showing reduced amygdala-PFC connectivity in adults is associated with impaired emotional responses [13,51], anxiety and depression [20,52,53], and increased aggressive behaviour and attention problems in children [40]. Increased amygdala reactivity is also seen in adolescents with MDD [54]. Medications and psychotherapy that treat depression increase functional coupling between the amygdala and the striatum, thalamus, right frontal and cingulate cortex [52], likely resulting in increased PFC inhibition and regulation of amygdala responsivity. With this in mind, it is important to note that although data consistently shows altered amygdala-prefrontal cortex connectivity associated with mood and anxiety disorders, it is unclear whether that dysfunction is hypo- or hyper-connectivity [46]. Our results support the theory of weaker top-down amygdala inhibition in children who experienced higher prenatal maternal depression and show subsequent behavioural alterations, and provide evidence of a structural basis for previously-observed deficits in functional connectivity.

Structural connectivity of the amygdala pathway mediated the relationship between maternal prenatal depressive symptoms and externalizing behaviour in boys. Higher externalizing symptoms have consistently been found in children whose mothers experienced prenatal depression [55]. Thus, this mediating relationship suggests that alterations to amygdala-PFC structural connectivity may be a mechanism via which maternal depression impacts children’s behaviour [8]. Furthermore, this altered connectivity may be a mechanism that increases children’s risk for later mental health difficulties, as higher externalizing symptoms in childhood are associated with higher risk of suicidality, substance use, and severity of psychopathology in adolescence and adulthood [29,56–58].

These data demonstrate an interesting pattern of sexual differentiation suggesting that boys are more vulnerable to maternal depressive symptoms. The mediation was only significant in males and relationships in the cingulum were stronger in boys. Males experience more prominent age-related gray matter volume decreases and white matter increases [59] and accelerated growth in limbic regions (including the cingulum) compared to females [60], perhaps underlying their increased susceptibility to environmental factors such as prenatal depression. Males are more susceptible to behavioural disturbances than females. In childhood, males have a higher prevalence of depression [61], and males but not females of postnatally depressed mothers have poorer cognitive function [62]. Male infants exposed to prenatal depression have lower motor skills, higher generalized anxiety, and sleep problems [6]. Thus, our data highlights that it may be an underlying brain vulnerability in males that predisposes them to these behaviour problems. Of note, previous data in adults showed no moderating effect of sex on the relationship between prenatal stress during the first half of pregnancy and FA of the cingulum [44,63]. However, the different exposure periods and age of the offspring may account for the contrasting results.

The mechanisms through which prenatal maternal depressive symptoms influence child brain structure are not well understood. Theories include dysregulation of the hypothalamic-pituitary-adrenal (HPA) axis and increased cortisol during pregnancy, genetic heritability of mood disorders, epigenetic modification of pre-existing genes, nutrition, and increased inflammation [64–66]. Maternal stress and depression during pregnancy increase cortisol and corticotropin releasing hormone (CRH), which initiates signalling in the HPA [67] and can cross the placenta [64]. Cortisol exposure peaks in the third trimester at 2-3 times non-pregnant values [67]. High levels of stress in women during the 28-32^nd^ weeks of pregnancy have been associated with reductions in uterine blood flow, suggesting that fetuses may experience periods of hypoxia [68]. Depression is associated with HPA axis dysregulation, with increased and prolonged responses to stress and higher circulating glucocorticoids [69], which activate receptors in the child’s brain, including the cingulate cortex and amygdala and induces epigenetic changes [70]. MDD is highly heritable [71], and pre-existing genetic vulnerability likely plays an important role in child vulnerability. Poor nutrition during pregnancy increases the risk of perinatal depression and poor child cognitive outcomes, and thus may also play a role [72,73]. Stress during pregnancy is also associated with increased inflammation and cytokine activity particularly in the third trimester [65], which may also impact child outcomes [74]. It is likely a combination of these biological factors, as well as psychosocial factors like socioeconomical status, education, and social support [66], that contribute to altered brain development and child outcomes.

There are some limitations to this study. First, due to the timing of recruitment, limited data was available on maternal first trimester depressive symptoms, and therefore this data was not analyzed. Another limitation is that although greater severity of EDS depressive symptom scores is associated with a diagnosis of MDD, the EDS itself is not diagnostic. Therefore, although we could grade severity of symptoms, we cannot say if these women had a clinical diagnosis of depression. Future studies with a larger sample size of first trimester scores will be better able to answer questions of how prenatal timing of exposure to maternal depression symptoms is related to child outcomes.

In conclusion, we show altered structural connectivity of the amygdala in children of mothers with higher prenatal depressive symptoms, and these alterations mediate behavioural dysfunction in boys. These data present early evidence of a structural basis for decreased top down inhibition of the amygdala resulting in altered behaviour in preschoolers, and may indicate a structural abnormality predisposing these children to develop affective disorders. Indeed, these data may provide an explanation for why children born to depressed mothers have a much higher risk of developing depression themselves [8]. Additionally, we provide evidence of sexual differentiation, suggesting that males have increased vulnerability to changes in early childhood. The clinical implications of these findings are a need to better recognize and address prenatal depression as an important factor in child health outcomes.

## Acknowledgements and Disclosures

This research was supported by the Canadian Institute of Health Research (CIHR) (funding Grant Nos. IHD-134090 and MOP-136797 and MOP-123535), Alberta Innovates – Health Solutions, a grant from the Alberta Children’s Hospital Foundation, and the National Institute of Environmental Health Sciences (Grant No. 5R21ES021295-03). R.E.H was supported by a Mach-Gaensslen Foundation award through the University of Calgary Cumming School of Medicine. J.E.R was supported by an Eyes High University of Calgary Postdoctoral Scholarship and the T. Chen Fong Postdoctoral Fellowship in Medical Imaging Science. M.N.G was supported by a University of Calgary Queen Elizabeth II Graduate Studentship award and funding provided through the Department of Pediatrics, University of Calgary to D.D. We thank the members of the Alberta Pregnancy Outcomes and Nutrition study team: Catherine J. Field, Rhonda C. Bell, Francois P. Bernier, Marja Cantell, Linda M. Casey, Misha Eliasziw, Anna Farmer, Lisa Gagnon, Laki Goodewardene, David W. Johnson, Libbe Kooistra, Brenda M.Y. Leung, Donna P. Manca, Jonathan W. Martin, Linda J. McCargar, Maeve O’Beirne, Victor J. Pop, and Nalini Singhal

